# Drug mechanism-of-action discovery through the integration of pharmacological and CRISPR screens

**DOI:** 10.1101/2020.01.14.905729

**Authors:** Emanuel Gonçalves, Aldo Segura-Cabrera, Clare Pacini, Gabriele Picco, Fiona M. Behan, Patricia Jaaks, Elizabeth A. Coker, Donny van der Meer, Andrew Barthorpe, Howard Lightfoot, GDSC Screening Team, Andrew R. Leach, James T. Lynch, Ben Sidders, Claire Crafter, Francesco Iorio, Stephen Fawell, Mathew J. Garnett

## Abstract

Low success rates during drug development are due in part to the difficulty of defining drug mechanism-of-action and molecular markers of therapeutic activity. Here, we integrated 199,219 drug sensitivity measurements for 397 unique anti-cancer drugs and genome-wide CRISPR loss-of-function screens in 484 cell lines to systematically investigate *in cellular* drug mechanism-of-action. We observed an enrichment for positive associations between drug sensitivity and knockout of their nominal targets, and by leveraging protein-protein networks we identified pathways that mediate drug response. This revealed an unappreciated role of mitochondrial E3 ubiquitin-protein ligase *MARCH5* in sensitivity to MCL1 inhibitors. We also estimated drug on-target and off-target activity, informing on specificity, potency and toxicity. Linking drug and gene dependency together with genomic datasets uncovered contexts in which molecular networks when perturbed mediate cancer cell loss-of-fitness, and thereby provide independent and orthogonal evidence of biomarkers for drug development. This study illustrates how integrating cell line drug sensitivity with CRISPR loss-of-function screens can elucidate mechanism-of-action to advance drug development.

## Introduction

Understanding drug mechanism-of-action and evaluating *in cellular* activity is challenging (Santos *et al*, 2017) and widespread target promiscuity contributes to low success rates during drug development (Klaeger *et al*, 2017). For target-based drug development, a detailed understanding of drug mechanism-of-action provides information about specificity and undesirable off-target activity which could lead to toxicity and reduced therapeutic window (Lin *et al*, 2019). Moreover, molecular biomarkers can be used to monitor drug activity and to identify contexts in which drugs are more effective as the basis for patient stratification during clinical development.

The cellular activity of a drug is influenced by multiple factors including the selectivity and affinity of the compound to its target(s) and the penetrance of target engagement on cellular phenotypes. An array of biochemical, biophysical, computational and cellular assays are currently used to investigate drug mechanism-of-action (Schenone *et al*, 2013). For example, protein kinase inhibitors are profiled *in vitro* for their specificity and potency against panels of purified recombinant protein kinases. While informative, this approach fails to recapitulate the native context of the full-length protein in cells which could influence drug activity, it does not identify non-kinase off-target effects, nor is it suitable to evaluate the selectivity of compounds to non-kinase targets. Existing *in cellular* based approaches include transcriptional profiling following drug treatment of cells, chemical proteomics approaches such as kinobeads to measure drug-protein interactions, and multiplexed imaging or flow-cytometry to measure multiple cellular parameters upon drug treatment (Subramanian *et al*, 2017; Li *et al*, 2017; Reinecke *et al*, 2019). Despite the utility of these different approaches, gaining a full picture of drug mechanism-of-action, particularly in cells, remains a challenge and new approaches would be beneficial.

Pharmacological screens (Barretina *et al*, 2012; Garnett *et al*, 2012; Iorio *et al*, 2016; Subramanian *et al*, 2017; Lee *et al*, 2018) have been used to profile the activity of hundreds of compounds in highly-annotated collections of cancer cell lines with the aim of identifying molecular markers of drug sensitivity to guide clinical development (Cook *et al*, 2014; Nelson *et al*, 2015). More recently, CRISPR-based gene-editing has enabled the evaluation of highly specific and penetrant gene-knockout effects on cell fitness genome-wide in hundreds of cancer cell lines (Jinek *et al*, 2012; Shalem *et al*, 2014; Hart *et al*, 2015; Meyers *et al*, 2017; Behan *et al*, 2019). This has provided rich functional resources to explore cancer vulnerabilities and new potential drug targets (Marcotte *et al*, 2016; Meyers *et al*, 2017; Tsherniak *et al*, 2017; Behan *et al*, 2019). Parallel integration of gene loss-of-function screens with drug response can be used to investigate drug mechanism-of-action (Deans *et al*, 2016; Subramanian *et al*, 2017; Jost & Weissman, 2018).

Here, we integrate recent genome-wide CRISPR-Cas9 loss-of-function screens with pharmacological data for 397 unique anti-cancer compounds in 484 cancer cell lines. We show that CRISPR-Cas9 datasets recapitulate drug targets, can provide insights into drug potency and selectivity, and define cellular networks underpinning drug sensitivity. This approach identified a link between mitochondrial ubiquitin ligase *MARCH5* in MCL1 inhibitors response, and specifically in breast cancer cell lines. Furthermore, we defined robust pharmacogenomic associations, represented by genetic biomarkers independently supported by drug response and gene fitness measurements. These identify genetic contexts associated with drug-pathway dependency and provide a more refined set of biomarkers. Taken together, we present here an approach to leverage pharmacological and CRISPR screening data to inform on drug *in cellular* mechanism-of-action to guide drug development.

## Results

### Cancer cell line drug sensitivity and gene fitness effects

We analysed datasets from a highly-annotated collection of 484 histologically diverse human cancer cell lines (Supplementary Table 1). These have been extensively genetically characterised and utilised for large-scale drug sensitivity testing and CRISPR-Cas9 whole-genome loss-of-function screens (Garnett *et al*, 2012; Iorio *et al*, 2016; Meyers *et al*, 2017; Picco *et al*, 2019; van der Meer *et al*, 2019). We expanded on published single agent drug sensitivity data (Garnett *et al*, 2012; Lynch *et al*, 2016; Iorio *et al*, 2016; Picco *et al*, 2019) to consider 199,219 IC50 values for 397 unique cancer drugs (480 drugs including duplicates, Supplementary Table 2). These encompassed FDA-approved cancer drugs, drugs in clinical development, and investigational compounds with multiple modes of action, including 24 chemotherapeutic agents and 367 small molecule inhibitors. Drugs considered in this study had a response in at least 3 cell lines (IC50 lower than half of the maximum screened concentration) and 86% of all possible drug/cell line IC50 measurements have been evaluated (Supplementary Figure 1a, Supplementary Table 3). Two experimental protocols were used to generate the drug sensitivity measurements, termed here as GDSC1 (Iorio *et al*, 2016) and GDSC2 (Picco *et al*, 2019), chronologically ordered (Supplementary Figure 1b). A principal component analysis (PCA) of IC50 values identified a screen specific batch effect associated with principal component (PC) 2 which explained 2.8% of the total variance (Supplementary Figure 1c). For this reason, despite the fact that compounds screened with both technologies showed good agreement (n=66, mean Pearson’s R=0.50), we analysed the measurements of the screens separately. Analysis of the drug response variation across cell lines revealed that PC 1 (28.7% variance captured) was significantly correlated with cell line growth rate (Pearson’s R=-0.51, p-value=1.2e-28), particularly for chemotherapy agents and growth inhibitors (Supplementary Figure 1d and 1e).

Cell fitness effects for 16,643 gene knockouts have been measured using genome-wide CRISPR-Cas9 screens at the Sanger and Broad Institutes (Meyers *et al*, 2017; Behan *et al*, 2019; DepMap, 2019) (Supplementary Table 4). The first PC across the cell lines (6.8% variance explained) separated the two institutes of origin (Supplementary Figure 2a), consistent with a comparative analysis performed on an overlapping set of cell lines (Dempster *et al*, 2019). Growth rate was less significantly associated with CRISPR knockout response (Supplementary Figure 2b and 2c).

### Gene knockout fitness effects correspond with drug targets

We began by investigating the extent to which drug sensitivity corresponded to CRISPR knock-out of drug targets. We systematically searched for associations between drug sensitivity and gene fitness effects across the 484 cell lines (Figure 1a). We expect this to capture a variety of relationships ranging from direct drug-target interactions to more complex associations arising from interactions with regulators of the drug target(s). We tested a total of 7,988,640 single-feature gene-drug associations using linear mixed regression models. Potential confounding effects such as growth rate, culture conditions, data source and sample structure were considered in the models. We identified 865 significant associations (FDR adjusted p-value < 10%, Supplementary Table 5) between drug response and gene fitness profiles (Figure 1b), termed hereafter as significant drug-gene pairs. For this analysis we were able to manually curate the nominal therapeutic target(s) for 94.7% (n=376) of the anti-cancer drugs (Supplementary Figure 3a and Supplementary Table 1).

**Figure 1.**
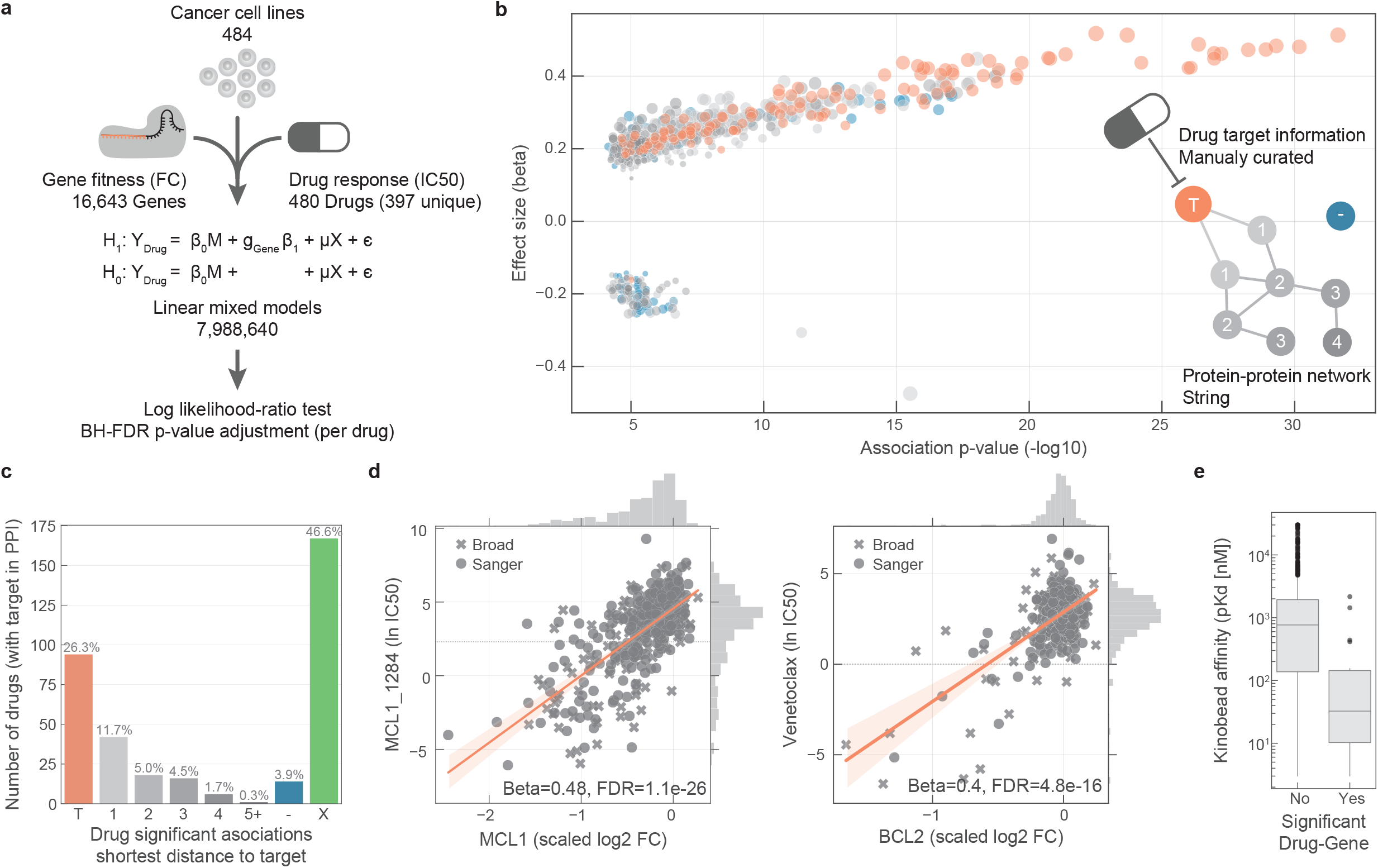
Integration of drug and gene dependencies in cancer cell lines. **a**, we used linear models to integrate drug sensitivity and gene fitness measurements. **b**, volcano plot showing the effect sizes and the p-value for statistically significant associations, Benjamini-Hochberg False Discovery Rate (FDR) adjusted p-value < 10%. Drug-gene associated pairs are coloured according to their shortest distance in a protein-protein interaction network of the gene to any of the nominal target of the drug. **c**, percentage of the 358 drugs with significant associations and their closest distance to the drug nominal targets. T represents drugs that have a significant association with at least 1 of their canonical targets and X are those which have no significant association. **d**, representative examples of the top drug response correlations with target gene fitness. MCL_1284 and Venetoclax are MCL1 and BCL2 selective inhibitors, respectively. Gene fitness log2 fold-changes (FC) are scaled by using previously defined sets of essential (median scaled log2 FC = −1) and non-essential (median scaled log2 FC = 0) genes. Drug response IC50 measurements are represented using the natural log (ln IC50). **e**, kinobead affinity is significantly higher (lower pKd) for compounds with a significant association with their target (n = 20, Mann-Whitney p-value=3.1e-07).

For 26% (n=94) of the 358 drugs with target annotation and for which the target was knocked-out with the CRISPR-Cas9 libraries, we identified significant drug-gene pairs with their putative targets (Figure 1c). For example, there were strong associations between MCL1 and BCL2 inhibitors and their knockouts (Figure 1d). Notably, drug-gene associations with the drug target had a skewed distribution towards positive effect sizes (Mann–Whitney U test p-value < 1.36e-105, Supplementary Figure 3b) and were among the strongest associations (Figure 1b). To investigate this further, we utilised independently acquired kinobead drug-protein affinity measurements for an overlapping 64 protein kinase inhibitors which were profiled for their specificity against 202 kinases (Klaeger *et al*, 2017). Drugs with significant associations with the target also had stronger affinity to their target in the kinobead assay, providing independent evidence that the strongest drug-gene associations are enriched for targets of the drugs (Figure 1e). Overall, we identified the nominal target of approximately one quarter of the drugs tested using orthogonal CRISPR gene fitness screens, and drug targets were amongst the most significant associations.

### Cellular networks underpinning drug response

The remaining 74% (n=264) of drugs were not significantly associated with the CRISPR loss-of-function measurements of their nominal targets (Figure 1c). We reasoned that superimposing the significant drug-gene pairs onto a protein interaction network may shed further insight into drug mechanism-of-action. We used a protein-protein interaction (PPI) network assembled from STRING database (Szklarczyk *et al*, 2017) (10,587 nodes and 205,251 interactions), and for the significant drug-genes pairs calculated the distances between the drug nominal targets and the associated gene-products. For 76 drugs no significant association with their target was identified, but instead had a significant association with their target’s first neighbour or a protein closely related in the network (1, 2 or 3 PPI interactions distance from any of the drug targets) (Figure 1b and c). Thus, 47.5% of the annotated compounds (n=170) had an association with either the target or a functionally-related protein.

The strongest drug-gene pair associations were between the drug and the canonical targets rather than components of the PPI network, and significance decreased (along with the number of associations) as the interaction distance increased (Figure 2a). To exclude that this observation is related to the topology of the network, we calculated the length of all the shortest paths between the drug targets and their associated genes and confirmed the enrichment of first and second neighbours in significant drug-gene associations (Supplementary Figure 3c). In comparison, cell line gene expression profiles are less powered to identify associations with the PPI neighbours of the drug target (Figure 2b; Supplementary Table 6). In particular, the number of drugs significantly associated with their targets substantially decreased (n=17) and significant associations were predominantly found with gene-products further away in the PPI network and close to the average length of all paths (lG = 3.9). As an example, MIEN1 gene expression is significantly correlated with multiple EGFR and ERBB2 inhibitors which can be explained, not by a functional relationship, but by genomic co-localisation with *ERBB2* on chromosome 17. Hence, CRISPR measurements are more powered than gene expression to identify drug functional interaction networks.

**Figure 2.**
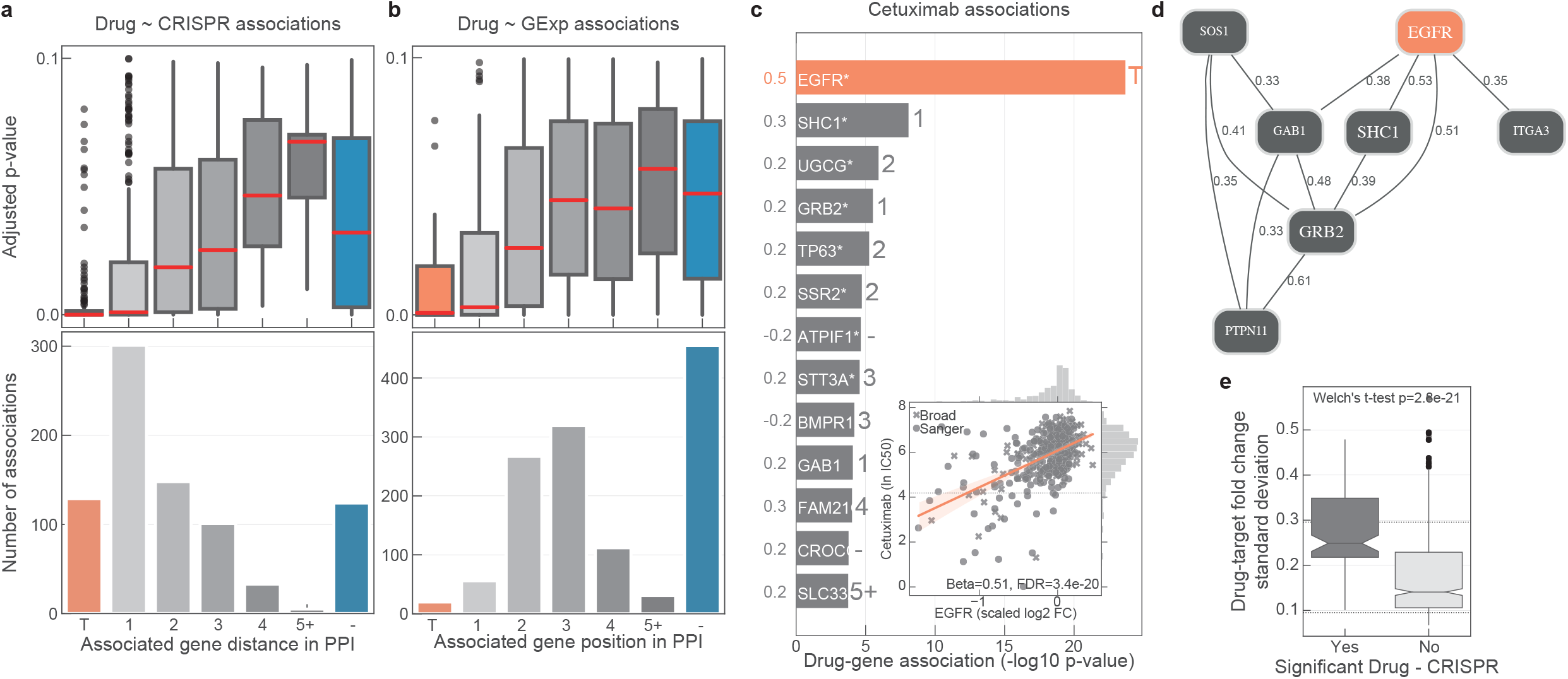
Drug response protein-protein networks. **a**, distribution of the FDR adjusted p-values (top) and count (bottom) of the significant drug-gene (CRISPR) associations according to their distance between the gene and corresponding drug targets in the protein-protein interaction network. **b**, similar to **a**, but instead gene expression (GExp) was tested to identify associations with drug response. **c**, representative example (cetuximab - EGFR inhibitor) of the associations and **d**, networks that can be obtained from the integrative analysis. Edges in the network are weighted with the Pearson correlation coefficient obtained between the fitness profiles of interacting nodes. **e**, drug-target associations stratified by statistical significance and plotted against the standard deviation of the drug-target CRISPR fold changes. Upper and lower dashed lines represent the standard deviations of essential and non-essential genes, respectively.

To investigate putative regulatory networks for drugs, we weighted the PPI network edges with the correlation between the fitness profiles of the two connected nodes and integrated the resulting weighted network with drug response associations. *EGFR* inhibitors are the most abundant drug class in our set, and we observed that multiple inhibitors (e.g. cetuximab) showed significant associations with *EGFR* and known pathway members, for example *SHC1* and *GRB2* (Zheng *et al*, 2013; Scaltriti & Baselga, 2006) (Figure 2c). Additionally, the weighted network shows pathway members that have strongly correlated fitness profiles, which are likely functionally related (Pan *et al*, 2018). For EGFR inhibitors these included tyrosine receptor kinases *NTRK3* and *MET*, and the protein phosphatase *PTPN11* (Wang *et al*, 2017; Pan *et al*, 2018) (Figure 2d). Drug-target tailored networks can be used to understand drug mechanism-of-action, and have the potential to identify resistance mechanisms and propose alternative targets in the network.

Despite our finding that we can illuminate drug functional networks, 46.6% (n=167) of the tested drugs had no significant drug-gene associations. This could in part be explained by lower variability in CRISPR fold change measurements for the target of these drugs (Figure 2e). For example, where genetic knockout induces strong uniform loss-of-fitness effects in contrast to incomplete target inhibition by a drug (Supplementary Figure 3d). Additionally, inhibition of a protein is intrinsically different than a knockout, as observed for PARP inhibitors whose activity is mediated through formation of cytotoxic PARP-DNA complexes, whereas as PARP knockout has little or no effect on cells (Gill *et al*, 2015; Murai & Pommier, 2015)(Supplementary Figure 3e). A lack of variability was much less pronounced in the drug sensitivity measurements since we only considered drugs which showed a minimal level of activity, i.e. IC50 lower than half of the maximum screened concentration (Supplementary Figure 3f). Drugs with no significant association were also approximately 3 times less likely to be associated with a genomic biomarker linked to sensitivity (Supplementary Figure 3g). Thus, the absence of an association between drug sensitivity and CRISPR loss-of-function effects could warrant further investigation into drug mechanism-of-action to understand possible underlying factors, such as low potency, alternative molecular mechanisms, or polypharmacology. Collectively, our network analysis demonstrates that CRISPR screens can provide functional insights into drug *in cellular* activity extending beyond the direct drug target into the associated functional network.

### Cancer drugs mechanism-of-action

Next, we set out to investigate in detail some of the strongest drug sensitivity and gene fitness associations (Supplementary Table 5). Strikingly, 46 of the top 50 strongly associated drugs have significant associations with their nominal target and with known functionally related genes (Figure 3). Some of the strongest associations were between MCL1 inhibitors and their target fitness effects (Figure 1d), including AZD5991 which is currently in clinical trials for treatment of hematologic cancers (Hird *et al*, 2017). Additionally, for several Insulin-Like Growth Factor 1 Receptor (*IGFR1*) inhibitors the association with the target was recapitulated. Moreover, significant associations with proprotein convertase *furin* were observed, supporting the known genetic association that *IGFR1* is a *furin* substrate, and increased levels of *furin* are associated with increased levels of processed IGFR1 and worse prognosis in several cancers (Thomas, 2002).

**Figure 3.**
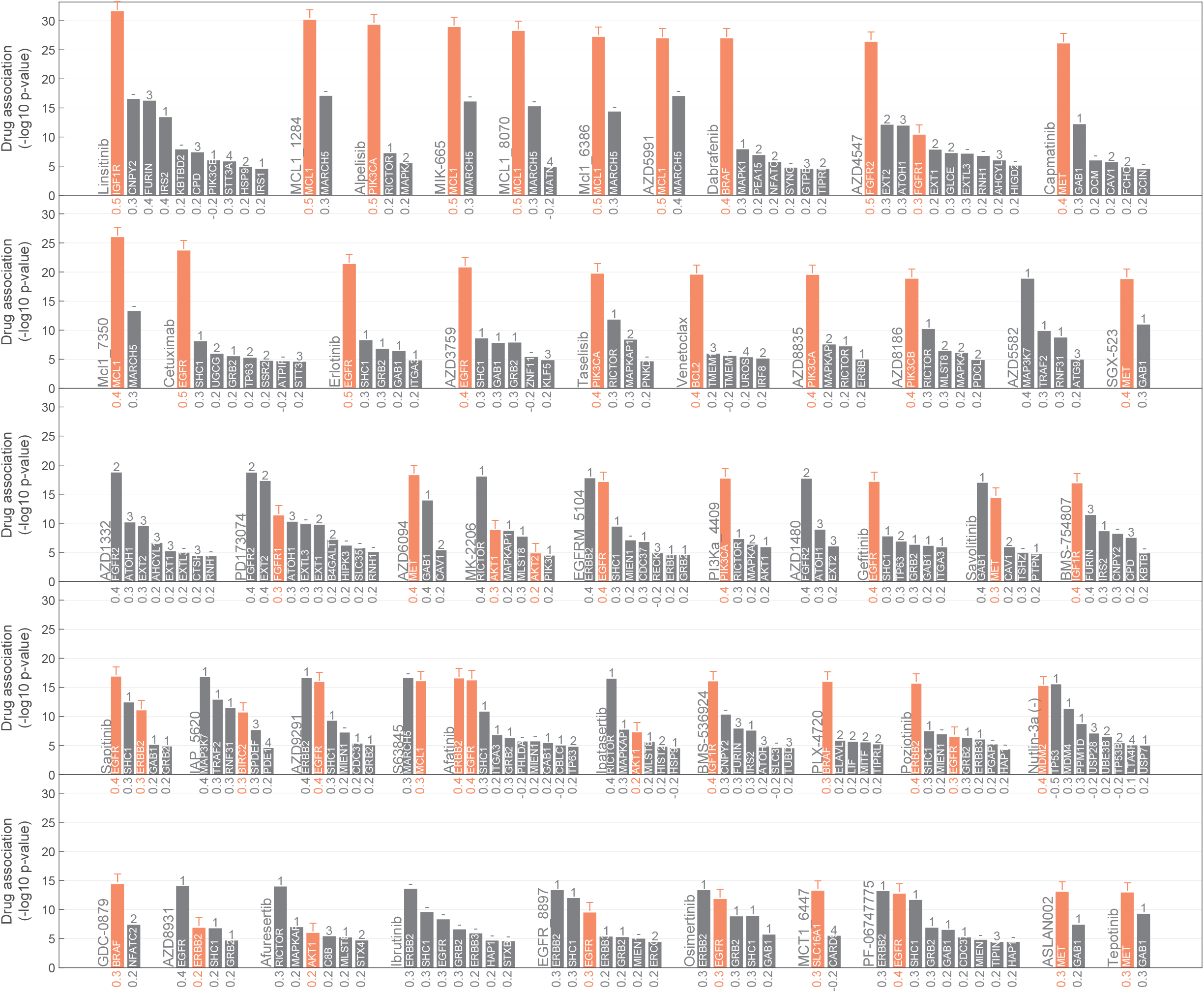
Top 50 most significantly associated drugs. Each bar plot group represents a unique drug where genes are ranked by statistical significance of their association. Effect sizes of the associations are reported under the bars along the x axis. Shortest distance (number of interactions) in a protein-protein interaction network between the gene and the drug nominal target(s) is represented on the top of the bars, where T and orange bar represent the target and represents no link was found.

Protein kinase inhibitors are an important class of cancer drugs (Santos *et al*, 2017). Because of the conserved structural features of the commonly targeted kinase domain, the clinical development of kinase inhibitors is hampered by poor selectivity, which consequently may lead to undesirable off-target activity (Klaeger *et al*, 2017). Furthermore, some kinases have multiple isoforms with non-redundant roles in tissues, as exemplified by the clinical development of PI3K inhibitors, and this has led to the development of isoform-selective inhibitors to reduce toxicity and increase the therapeutic window (Thorpe *et al*, 2015). Interestingly, several PI3K inhibitors had strong associations with only one gene encoding a single isoform (Figure 3), this together with the increased kinobead binding affinity of significant associations (Figure 1e), suggests these are isoform selective compounds. For example, alpelisib (Figure 3 first row) was associated with *PIK3CA*, consistent with its development as an alpha-isoform selective compound (Thorpe *et al*, 2015), whereas AZD8186 (Figure 3 second row) was only associated with *PIK3CB* confirming its beta-selectivity. Conversely, two *pan-*PI3K inhibitors (buparlisib and omipalisib) displayed no significant association with any *PI3K* isoform (Supplementary Table 5), consistent with less isoform specificity and potential polypharmacology. Interestingly, MTOR and pan-PI3K inhibitor, dactolisib, had significant associations with *RPTOR* and *MTOR* but none with *PI3K* isoforms (Supplementary Table 5), consistent with recently reported greater specificity for inhibition of the MTOR complex (Reinecke *et al*, 2019). Similarly, we observed that selective EGFR inhibitors cetuximab, erlotinib and gefitinib (Figure 3 second and third rows) were associated with *EGFR* but not *ERBB2*, whereas afatinib, poziotinib and sapatinib (AZD8931) (Figure 3 fourth and fifth rows) were all associated with both *EGFR* and *ERBB2*. Furthermore, we observed isoform selectivity of different FGFR inhibitors.

Our analysis can also provide insights into possible off-target activity of drugs. Unsupervised clustering of the drug-gene associations effect sizes (betas) revealed classes of inhibitors with similar targets and mechanism-of-action (Supplementary Figure 3h). Of note, BTK inhibitor, ibrutinib, clustered with EGFR inhibitors and displayed significant associations with EGFR and ERBB2 gene fitness. This is consistent with recent findings that brutinib covalently bind and inhibit EGFR (Lee *et al*, 2018), and is also supported by kinobead measurements (Klaeger *et al*, 2017). Additionally, 24 compounds have significant associations with genes identified as core fitness (Behan *et al*, 2019) across multiple cancer types, indicating an increased risk of cellular toxicity. Out of these, two compounds, PD0166285 and CCT244747, have significant associations with their nominal target (*PKMYT1* and *CHEK1*/*WEE1*) and the remaining compounds (n=22) are correlated with proteins closely connected in the PPI network.

### A functional link between *MARCH5* and MCL1 inhibitor sensitivity

Seven out of nine inhibitors of the anti-apoptotic *BCL2* family member myeloid cell leukemia 1 (*MCL1*) were strongly and nearly exclusively associated with their putative target, suggesting these are potent and specific compounds in cells (Figure 4a). *MCL1* is frequently amplified in human cancers (Beroukhim *et al*, 2010) and associated with chemotherapeutic resistance and relapse (Wuillème-Toumi *et al*, 2005; Wei *et al*, 2006). MCL1 is a negative regulator of the mitochondrial apoptotic pathway, regulating *BAX/BAK1* which co-localise with *Drp1/Fis1* in the mitochondria outer membrane and control mitochondrial fragmentation and cytochrome c release, both of which are important for inducing apoptosis (Youle & Karbowski, 2005; Mojsa *et al*, 2014; Morciano *et al*, 2016). Interestingly, knockout of a key regulator of mitochondrial fission, mitochondrial E3 ubiquitin-protein ligase *MARCH5* (Karbowski et al, 2007), is significantly associated with MCL1 inhibitors sensitivity (Supplementary Figure 4a), and positively correlated with MCL1 gene fitness, suggesting a functional relationship (Figure 4b). A recent study confirmed a synthetic-lethal interaction between *MARCH5* and well know *MCL1* negative regulator *BCL2L1* using dropout screens in isogenic cancer cell lines (DeWeirdt *et al*, 2019). Correlation between *MCL1* and *MARCH5* fitness profiles shows that cell lines dependent on *MARCH5* are also dependent on *MCL1*, while the inverse is not necessarily true with a subgroup of cell lines dependent on *MCL1* but not on *MARCH5*. Cell lines independently dependent on both gene-products have increased sensitivity to MCL1 inhibitors (Supplementary Figure 4b). This is particularly marked in breast carcinoma cancer cell lines, with *MCL1* and *MARCH5* dependent cells having similar sensitivity to hematologic cancer cell lines (acute myeloid leukemia), where MCL1 inhibitors are in clinical development (Figure 4c).

**Figure 4.**
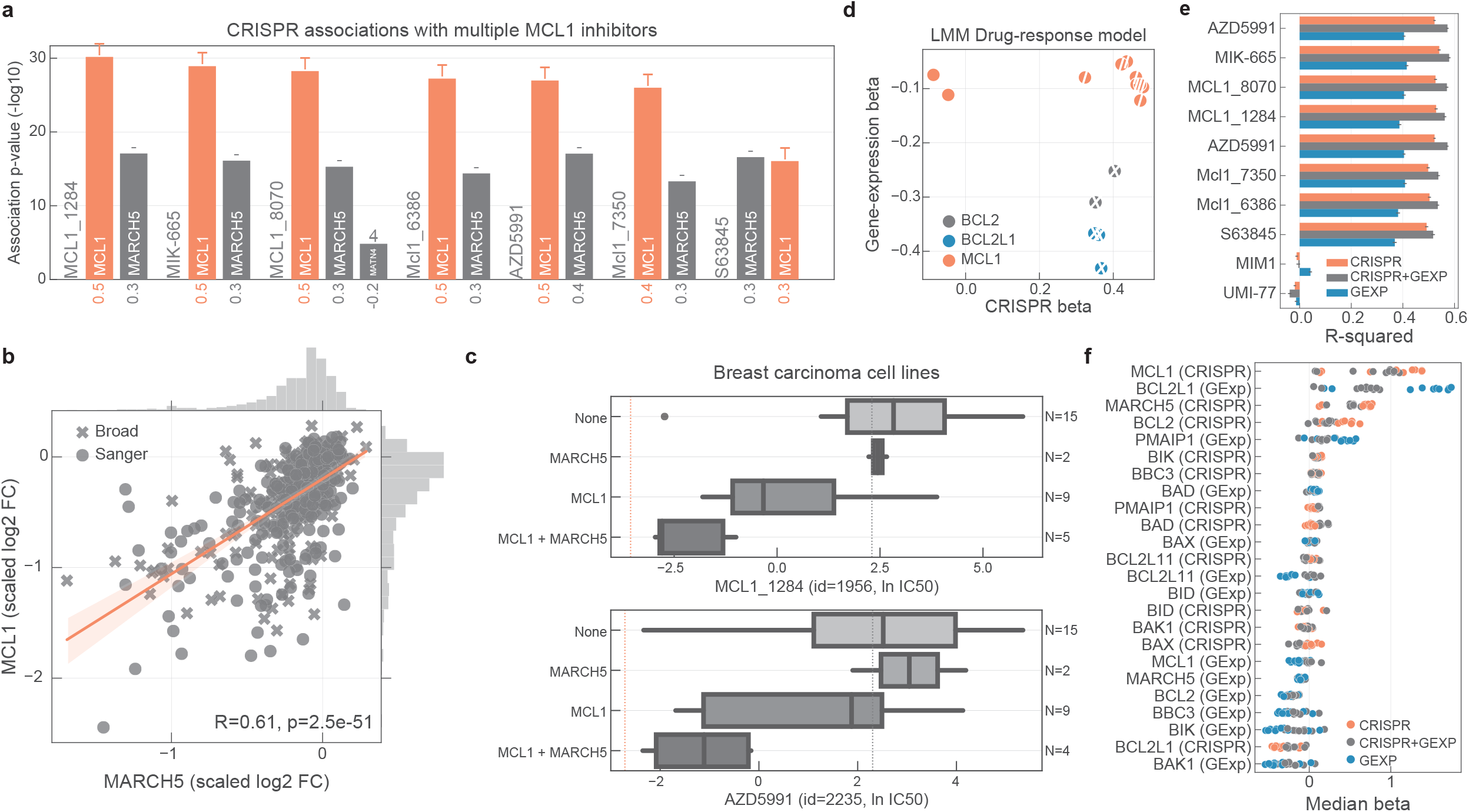
MCL1 inhibitors associations. **a**, significant associations with MCL1 inhibitors (7 out of 9 included in the screen). **b**, association between the gene fitness profiles of MCL1 and MARCH5. **c**, stratification of the MCL1 inhibitor sensitivity according to the essentiality profile of MCL1 and MARCH5, where MCL1 + MARCH5 represents a cell line that is independently dependent on both genes. Dashed orange line (left) represents the mean IC50 in acute myeloid leukemia cell lines. Grey dashed line (right) represents the maximum concentration used in the dosage response curve. **d**, BCL2, BCL2L1 and MCL1 inhibitors and the respective association with their targets, on the x axis with CRISPR gene fitness and on the y axis with gene expression. The statistical significance of the association is represented with a backward slash for CRISPR and forward slash for GEXP. **e**, regularised multilinear regression to predict drug response of all MCL1 inhibitors using gene expression, fitness or both of known regulators of the BCL2 family and MARCH5. Predictive performance is estimated using R2 metric represented in the x axis. **f**, effect size of each feature used in each MCL1 inhibitor model.

We investigated the potential molecular mechanisms underlying MCL1 inhibitors response. MCL1 copy number and gene expression alone are not a good predictor of MCL1 inhibitors sensitivity (Figure 4d, Supplementary Figure 4c). This is in contrast to BCL2 and BCL2L1 inhibitors, where their target gene expression is significantly correlated with drug sensitivity (Figure 4d). Next, we used multilinear regression models to predict sensitivity to each MCL1 inhibitor using gene fitness and/or gene expression of known regulators of *MCL1* (e.g. *BCL2, BCL2L1, BAX*) (Czabotar *et al*, 2014) and *MARCH5*. For two MCL1 inhibitors, MIM1 and UMI-77, the trained models performed poorly likely due to lack of *in cellular* activity of these compounds. For the remaining seven MCL1 inhibitors, drug response was well predicted (CRISPR+GEXP mean R-squared=0.55). Models trained with only CRISPR displayed overall better predictions compared to models only trained with gene expression, and models trained with both data types out-performed all others (Figure 4e). As expected, MCL1 fitness-effect was the most predictive feature, followed by BCL2L1 expression and MARCH5 essentiality (Figure 4f). No genomic feature, mutation or copy number alterations correlated significantly with MCL1 inhibitors response, including *MCL1* amplifications (Supplementary Figure 4c), likely a consequence of the strong post-transcriptional regulation and short half-life of MCL1.

Altogether, we highlight a functional link between *MARCH5* and MCL1 inhibitors sensitivity. With further investigation, this could shed light on MCL1 inhibitor mechanism and the development of stratification approaches in solid tumours, such as breast carcinomas.

### Robust molecular markers of drug sensitivity networks

The identification of molecular biomarkers of drug sensitivity is fundamental to guide clinical drug development. We hypothesized that molecular biomarkers independently linked with both drug response and gene fitness would be of particularly high value – termed robust pharmacogenomic biomarkers. To identify these, we used the set of significant drug-gene pairs (n=865) and we searched independently for significant associations between each measurement type in each pair (drug response or gene fitness) and 519 genomic (mutations and copy number alterations) and 15,368 gene expression features (Figure 5a, Supplementary Figure 5a) (Garnett *et al*, 2012; Iorio *et al*, 2016; Garcia-Alonso *et al*, 2018). This analysis recapitulated established genomic and expression biomarkers of either drug sensitivity or gene fitness effects in cancer cells (Supplementary Figure 5b and c). A total of 224 and 679 robust pharmacogenomic associations were identified with genomic (Supplementary Table 7) and gene expression features (Supplementary Table 8), respectively. Overall, 30.6% (265 of 865) of drug-gene pairs have at least one robust molecular marker that correlated significantly with both drug response and gene fitness (Figure 5b). The number of robust biomarkers was smaller than the number of biomarkers associated with only one type of measurement, likely due to the stringent requirement for an association with both drug sensitivity and gene fitness effects.

**Figure 5.**
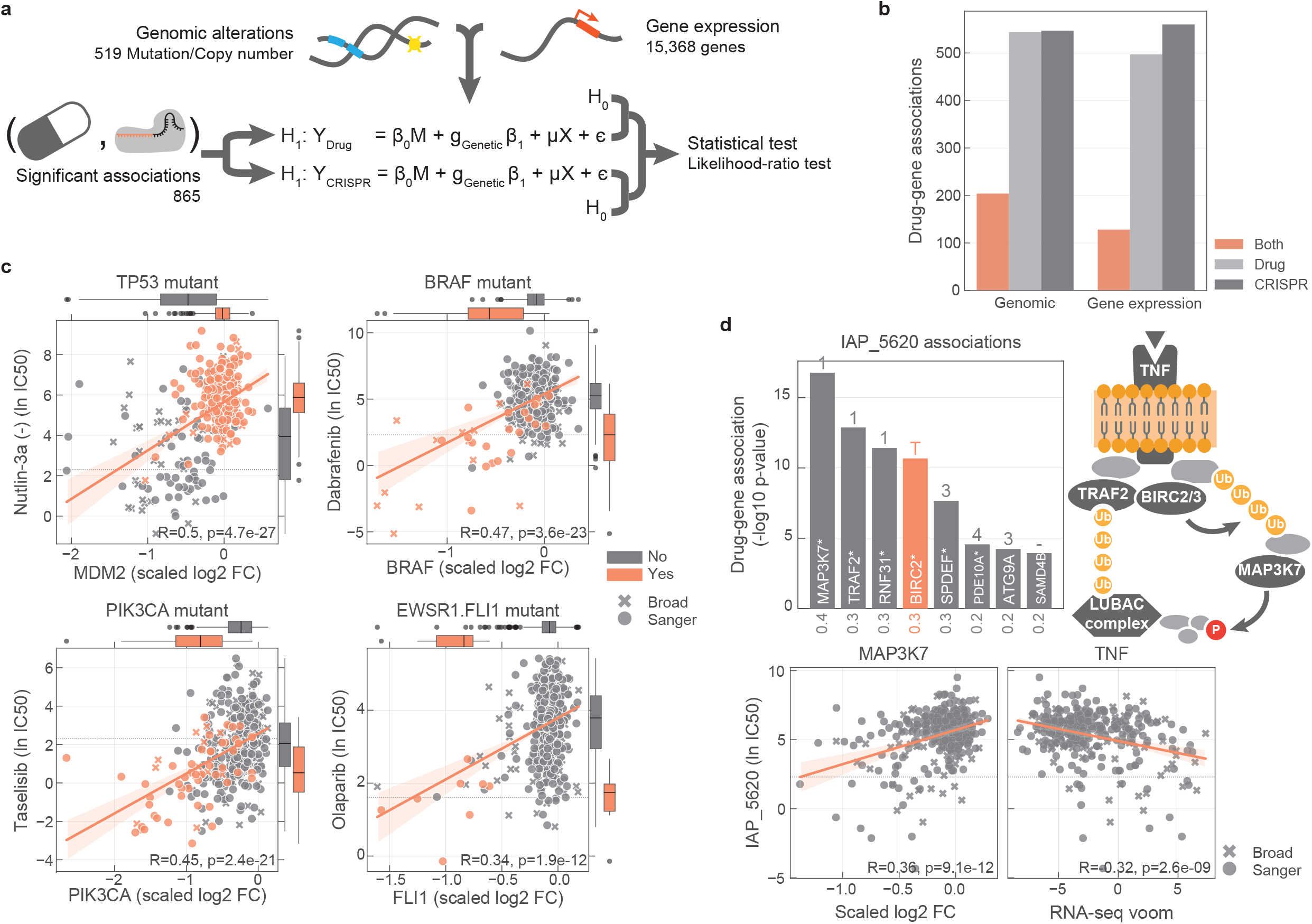
Robust pharmacological associations. **a**, diagram representing how genomic and gene expression data-sets are integrated to identify significant associations with drug-gene pairs that were previously found to be significantly correlated. **b**, number of drug-gene pairs with at least one significant association with drug response, gene fitness or both, considering either genomic or gene expression profiles. **c**, canonical examples of robust pharmacological associations. **d**, representative example of a BIRC2/BIRC3 inhibitor, IAP_5620, showing the significant associations with CRISPR gene fitness profiles and their location in a representation of the TNF pathway.

From the subset of 129 drug-gene pair associations that were linked by the drug target, 50.4% (n=65) had one or more robust pharmacogenomic associations (Supplementary Figure 5d). Most of these were established dependencies of cancer cells, including: Nutlin-3a sensitivity associated with TP53 mutation status; BRAF and PIK3CA mutation induced CRISPR dependency; olaparib sensitivity mediated by the presence of EWSR1-FLI1 fusion, also recapitulated by FLI1 essentiality profile; MCL1 inhibitors biomarker association with BCL2L1, and nutlin-3a with BAX expression (Figure 5c and Supplementary Figure 5e and 5f). Similarly, of the 413 significant gene-drug pairs closely related within the PPI network (<=3 interactions from the drug target), we identified robust pharmacogenomic associations for 29.5% (n=122) (Supplementary Figure 5d), enabling the discovery of cellular contexts where drug response networks are important. For example, we identified increased tumour necrosis factor (TNF) expression as a robust pharmacogenomic marker for drugs targeting the downstream cellular inhibitor of apoptosis (cIAP) proteins BIRC2 and BIRC3 (e.g. IAP_5620), and based on CRISPR dependency data, for multiple members of the cIAP pathway, including BIRC2, MAP3K7 and RNF31 (Beug et al, 2012) (Figure 5d).

## Discussion

Understanding drug mechanism-of-action and the biological pathways underpinning drug response is an important step in preclinical studies. Here, we demonstrate how the integration of drug sensitivity and CRISPR-Cas9 gene fitness data can be used to inform on multiple aspects of drug mechanisms in cells, including drug specificity and potency. Our analysis recapitulated drug targets for approximately a quarter of drugs tested and for approximately another quarter revealed associations enriched for proteins closely related with the drug target. Critically, the strength of these associations reflects specificity and polypharmacology of the cancer drugs. Furthermore, these associations define networks of protein interactions that are functionally related with drug targets and underpin drug response. This revealed a previously unappreciated interaction between *MARCH5* and MCL1 inhibitors, with potential utility to derive predictive models of MCL1 inhibitor response across multiple cancer types, and particularly in solid tumours such as breast carcinomas. Robust pharmacogenomic biomarkers leveraged both datasets to provide refined biomarkers that are correlated with both drug response and biological networks. Interestingly, the networks we have defined can provide alternative targets that are functionally related with the drug target and mediate similar effects on cell fitness, potentially providing strategies for combination therapies to limit therapy resistance.

Pre-clinical biomarker development is an important step in drug discovery and is associated with increased success rates during clinical development (Nelson *et al*, 2015). Traditionally this has been performed by building predictive models of drug response using mutation, copy number and gene expression (Iorio *et al*, 2016; Tsherniak *et al*, 2017). Here we extended this approach, and propose what we term as robust pharmacogenomic association - a drug response and gene fitness pair that are significantly correlated and are also both significantly related to the same molecular biomarker. This approach gives greater confidence in molecular biomarkers identified, since they are recapitulated using data from two orthogonal assays and provides markers at the level of the network. In addition, by focusing only on drugs involved in significant gene-drug pairs, we enrich for drugs most likely to have greater specificity, and thereby better enabling biomarker discovery.

Nearly half of the drugs did not have a significant association with gene fitness effects and may warrant further investigation. Possible explanations for this include: (i) drug polypharmacology which is difficult to deconvolute using single gene knockout data; (ii) intrinsic difference between protein inhibition and knockout; (iii) a dosage dependent response leading to incomplete inhibition of the drug target; (iv) functional redundancy between protein isoforms resulting in less penetrant effects with gene knockout; and (v) limitations of the sgRNA efficacy across the cancer cell lines. We expect that some of these issues can be addressed by expanding this analysis to integrate other types of functional genomic screens, such as CRISPR inhibition, which might mimic drug inhibition more closely.

This study extends previous efforts, and utilises new CRISPR loss-of-function datasets, to study drug mechanism-of-action in cells with unparalleled scale and precision. We anticipate this approach to be useful for many compounds, and could become a routine step during drug development. In particular, it is likely to have utility during the hit-to-lead optimisation stage of drug development to select lead chemical series and compounds with optimal potency and selectivity. The utility of this approach is likely to expand as the availability of CRISPR knock-out screening data, and other datasets such as CRISPR activation and inhibition, increases across ever larger collections of highly-annotated cancer cell models. In conclusion, this study illustrates a new approach for investigating *in cellular* drug mechanism-of-action that can be applied to multiple critical aspects of drug development.

## Materials and Methods

### 1. Cancer cell lines panel

The 484 cancer cell lines used in this manuscript have been compiled from publicly available repositories as well as private collections and maintained following the supplier guidelines. STR and SNP fingerprints were used to ensure cell lines selected were genetically unique and matched those in public repositories (http://cancer.sanger.ac.uk/cell_lines/download). Detailed cell line model information is available through Cell Model Passports database (https://cellmodelpassports.sanger.ac.uk/) (van der Meer *et al*, 2019). Cell lines growth rate is represented as the ratio between the mean of the untreated negative controls measured at day 1 (time of drug treatment) and the mean of the DMSO treated negative controls at day 4 (72 hours post drug treatment).

### 2. High-throughput drug sensitivity

Experimental details of both GDSC1 and GDSC2 screens can be found in the Genomics of Drug Sensitivity in Cancer (GDSC) project (www.cancerRxgene.org) (Yang *et al*, 2013). Cell viability and dose response curve fitting models were previously described in detail (Iorio *et al*, 2016; Vis *et al*, 2016). Maximum screened drug concentration (μM) are provided in Supplementary Table 1. Each compound was measured on average across 393 cell lines rendering a nearly complete matrix with only 14.2% missing values. All considered compounds displayed an IC50 lower than half of the maximum screened concentration in at least 3 cell lines. This filter ensures the compounds display an informative profile in at least a small subset of the cell lines. Drug nominal oncology target annotation was manually curated from literature (Supplementary Table 1).

### 3. Genome-wide CRISPR-Cas9 dropout screens

The CRISPR-Cas9 screens for the 484 cancer cell lines considered in this study (Supplementary Table 2) were assembled from two distinct projects, 320 were generated as part of Sanger DepMap Project Score (Behan *et al*, 2019) and 164 from the Broad DepMap version 19Q3 (Meyers *et al*, 2017; DepMap, 2019). Only cell lines that passed quality control filtering similarly to Behan et al. (2019) and with matched drug response measurements were considered. Different CRISPR-Cas9 sgRNA libraries were used in each project (Koike-Yusa *et al*, 2014; Doench *et al*, 2016; Tzelepis *et al*, 2016). Consequently, library-specific effects were present (Dempster *et al*, 2019) (Supplementary Figure 2a) which hampers averaging of cell lines that were screened in both data-sets. Thus, for the overlapping cell lines only data from Sanger DepMap Project Score was used. This also minimises potential cell line specific differences, for example due to genetic drift (Ben-David *et al*, 2018), and thereby increasing concordance with the drug response data-set also generated at the Wellcome Sanger Institute. Fold changes (log2) were estimated comparing samples with the respective control plasmid. As copy number profiles were not available for all of the cell lines, gene-independent deleterious effects induced by copy number amplifications in CRISPR-Cas9 screens (Aguirre *et al*, 2016; Munoz *et al*, 2016; Gonçalves *et al*, 2019) were corrected on a per sample basis using the unsupervised method CRISPRcleanR (Iorio *et al*, 2018). Replicates were mean averaged and gene level fold changes were estimated by taking the mean of all the mapping sgRNAs. Gene level fold changes were quantile normalised per sample and then median scaled using previously defined lists of cancer cell lines essential and non-essential genes (Hart *et al*, 2015), thus essential genes have a median log2 fold change of −1 and non-essential genes a median log2 fold change of 0. Only overlapping genes between the two libraries were considered, thus generating a full matrix of 16,643 genes across the 484 cell lines. A cell line was considered dependent on a gene if the knockout had a log2 fold change of at least 50% of that expected of essential genes (scaled log2 fold change < −0.5).

### 4. PCA of drug sensitivity and gene fitness

Principal component analysis (PCA) was performed using scikit-learn (v0.21.2) (Pedregosa *et al*, 2011) using sklearn.decomposition.PCA with default parameters and the number of components (n_components) set to 10. For the drug response data-set, and only for the PCA analysis, missing values of each drug were imputed using the drug mean IC50 response across the rest of the cell lines. Imputation was not required for the CRISPR-Cas9 data-set since the matrix had no missing values.

### 5. Drug response linear mixed model associations

Associations between drug response and gene fitness scores were performed using an efficient implementation of mixed-effect linear models available in the LIMIX Python module (v3.0.3) (Lippert *et al*, 2014; Casale *et al*, 2017). We considered the following covariates in the model: (i) binary variables indicating the institute of origin of the cell line CRISPR-Cas9 screen; (ii) principal component 1 of the drug response data-set which is a correlative of cell lines growth rate; and (iii) growing conditions (adherent, suspension or semi-adherent) represented as binary variables. Additionally, gene fitness similarity matrix of the samples is considered as random effects in the model to account for potential sample structure. Taken together, we fitted the following mixed linear regression model for each drug-gene pair:

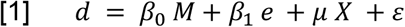

Where, *d* represents a vector of the drug response IC50 values across the cell lines; *M* is the matrix of covariates and *β*_0_ is the vector of effect sizes; e is the vector of gene CRISPR-Cas9 log2 fold changes and *β*_1_ the effect size; *X* the similarity matrix based on the CRISPR-Cas9 gene fitness measurements; *μ* is the random effects; *ε* is the general noise term. For each drug, cell lines with missing values were dropped from the fit.

We statistically assessed the significance of each association by performing likelihood ratio tests between the alternative model 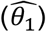 and the null model which excludes the gene CRISPR gene fitness scores vector *e* and its parameter 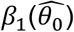. The parameter inference is performed using maximum likelihood estimation:

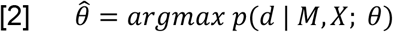

And the p-value of the association is defined by:

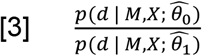

We tested all the single-feature pairwise associations between the 480 compounds and the 16,643 genes, making a total of 7,988,640 tested associations. P-value adjustment for multiple testing was performed per drug using the Benjamini-Hochberg False Discovery Rate (FDR). Contrary to performing the adjustment across all tests, per drug correction has the following benefits: (i) associations assembled from the different screening platforms (GDSC1 and GDSC2) are kept separate hence not biasing for measurement type; and (ii) drugs with stronger responses across larger subsets of cancer cell lines, for example Nutlin-3a response across TP53 wild-type cell lines, display stronger associations than most drugs, thus correcting across all drugs would retain more associations from these drugs at a specific error rate, i.e. 10%, compared to the rest.

### 6. Protein-protein interaction network

We assembled from STRING database (Szklarczyk *et al*, 2017) a high confidence undirected protein-protein interaction network. We only consider interactions with a combined confidence score higher than 900. Nodes’ STRING identifiers were converted to HUGO gene symbols, nodes not mapping or with multiple mappings were removed. Using *igraph* Python wrapper (Csardi & Nepusz, 2006) the network was simplified by removing unconnected nodes, self-loops and duplicated edges, leaving a total of 10,587 nodes and 205,251 interactions. A weighted version of the network was also assembled by correlating the gene fitness profiles of the connected nodes. Network nodes, and corresponding edges, that were not covered by the CRISPR-Cas9 screens were removed, making a total of 9,595 nodes and 172,584 weighted interactions.

### 7. Robust pharmacogenomic associations

Robust pharmacological associations were estimated similarly to the previous associations, but in this case only drug-gene pairs that are significantly correlated were considered to test associations with the genomic features (binarised copy number and mutation status (Iorio *et al*, 2016)) and gene expression profiles (RNA-seq voom (Law *et al*, 2014) transformed RPKMs (Garcia-Alonso *et al*, 2018)). A robust pharmacogenomic association is defined as: (i) a drug-gene pair whose drug sensitivity and gene fitness is significantly correlated, and (ii) genomic alteration or gene expression profile is significantly correlated with both drug response and gene fitness. Log-ratio test p-values are independently estimated for drug response and gene fitness measurements and corrected per drug-gene. Drug-gene pairs associated to a genomic or gene expression feature with an FDR lower than 10% are called robust pharmacogenomic associations (Supplementary Tables 7 and 8).

### 8. Predictive models of drug response of MCL1 inhibitors

L2-regularised linear regression models to predict MCL1 inhibitors drug response were trained using gene fitness, gene expression measurements or both of canonical regulators of *MCL1*, namely *MARCH5, MCL1, BCL2, BCL2L1, BCL2L11, PMAIP1, BAX, BAK1, BBC3, BID, BIK*, BAD. For the 9 MCL1 inhibitors considered in this study predictive models of drug response measurements were trained using Ridge regressions with an internal cross-validation optimisation of the regularization parameter, implemented in Sklearn with RidgeCV class (Pedregosa *et al*, 2011). Additionally, drug response measurements are split randomly 1,000 times, where 70% of the measurements are for training the model and 30% are left out as a test set. Model’s performance is quantified using the R2 metric on the test set, comparing the predicted versus the observed drug response measurements.

### 9. Code and data availability

Source code, analysis reports and Jupyter notebooks are publicly available in GitHub project https://github.com/EmanuelGoncalves/dtrace. Drug response and gene fitness CRISPR-Cas9 data-sets used in this analysis are available in the supplementary tables and accessible through figshare on https://doi.org/10.6084/m9.figshare.10338413.v1.

## Supplementary Figures

**Supplementary Figure 1.**
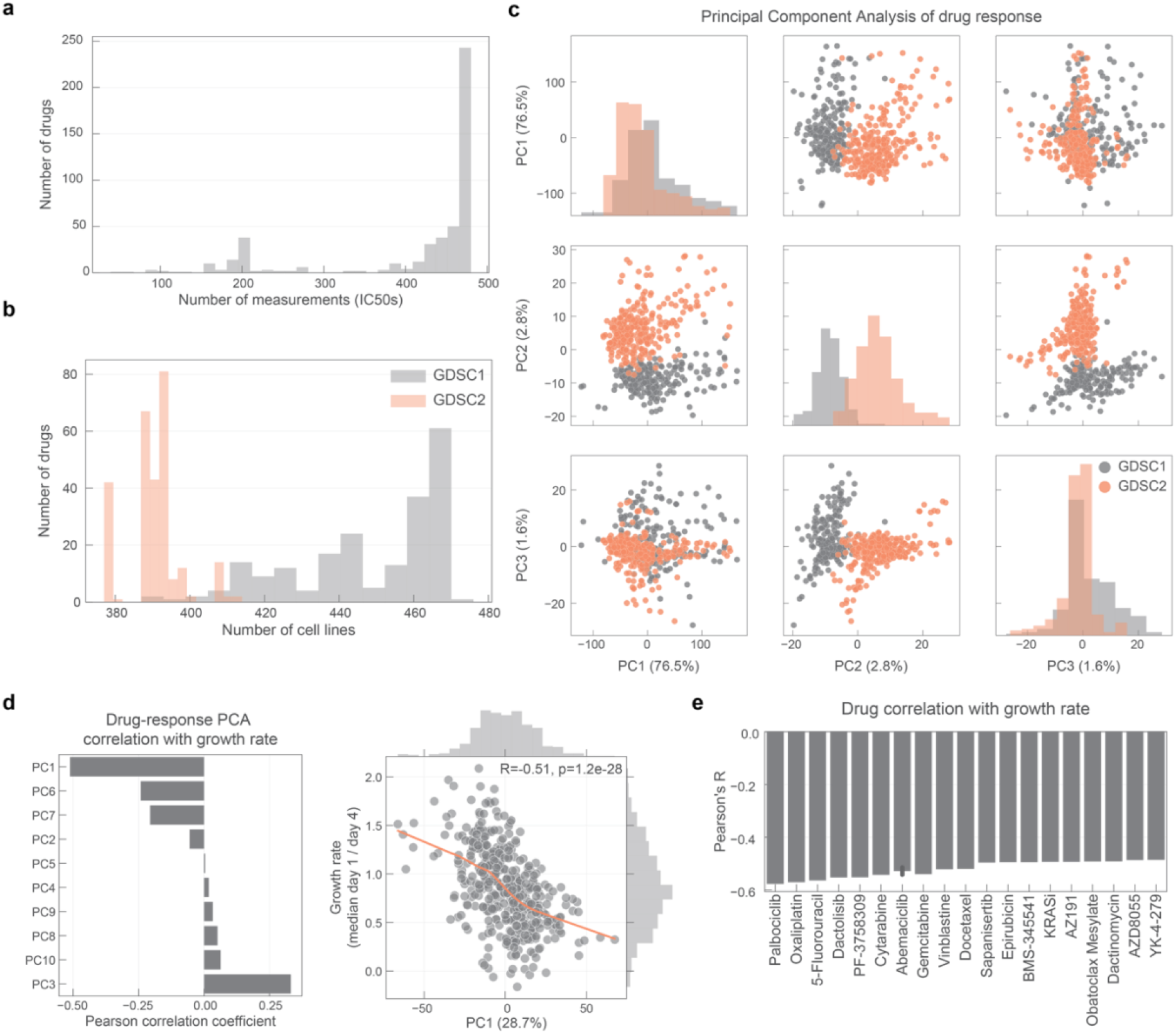
Overview of the drug sensitivity datasets. **a**, histogram of the number of IC50 values measured per drug. **b**, number of drugs measured per cell line in each pharmacological dataset. **c**, PCA analysis of the drug response measurements separated by the screen type. **d**, Pearson correlation coefficient between each principal component (PC) and cell lines growth rate. **e**, top absolutely correlated drugs with growth rate.

**Supplementary Figure 2.**
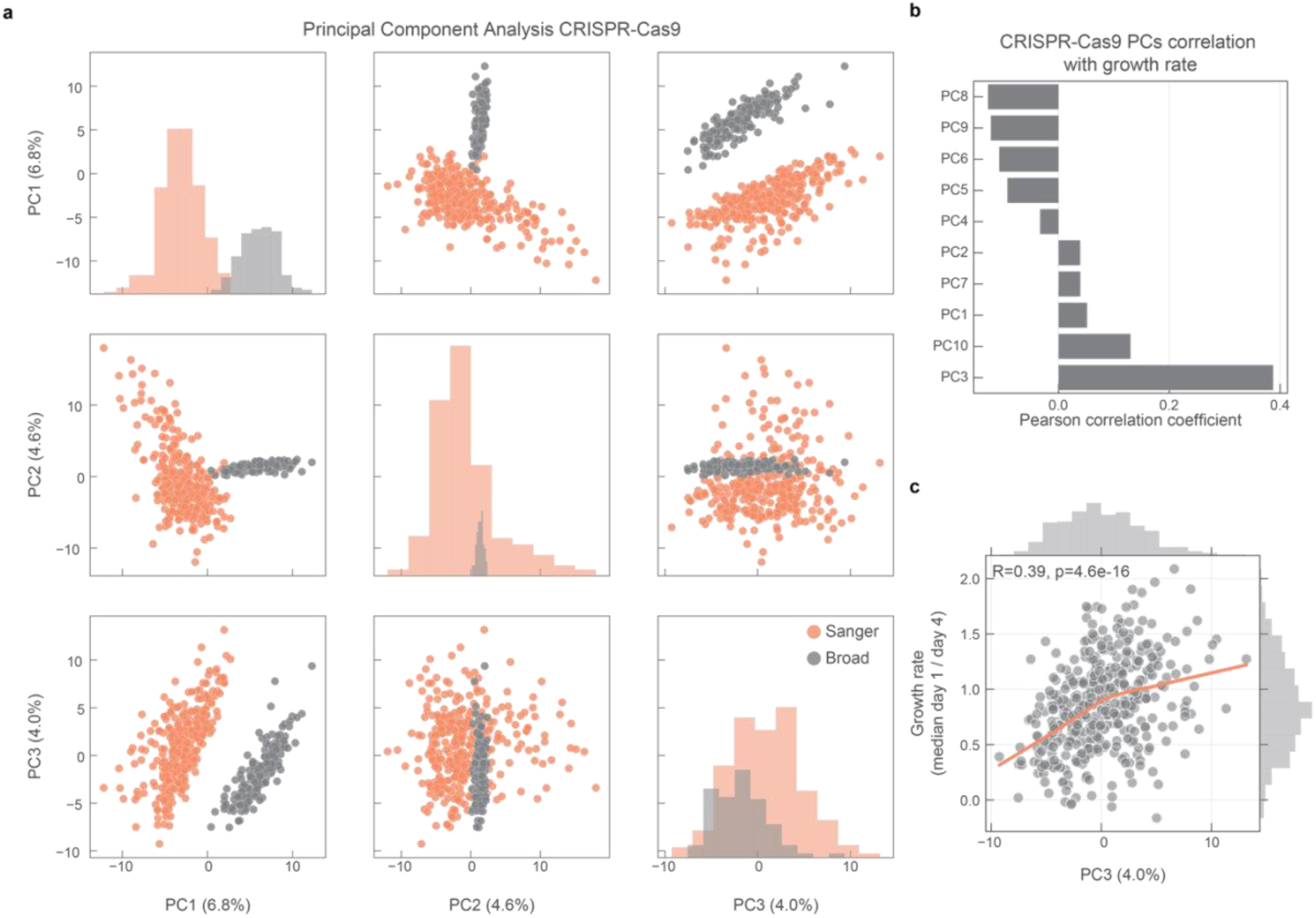
Overview of the CRISPR-Cas9 datasets. **a**, PCA analysis of the samples in the CRISPR-Cas9 screens, samples institute of origin is highlighted. **b**, correlation coefficients between all top 10 PCs and growth rate. **c**, correlation between cell lines growth rate and PC3 (Pearson correlation coefficient reported in the top left).

**Supplementary Figure 3.**
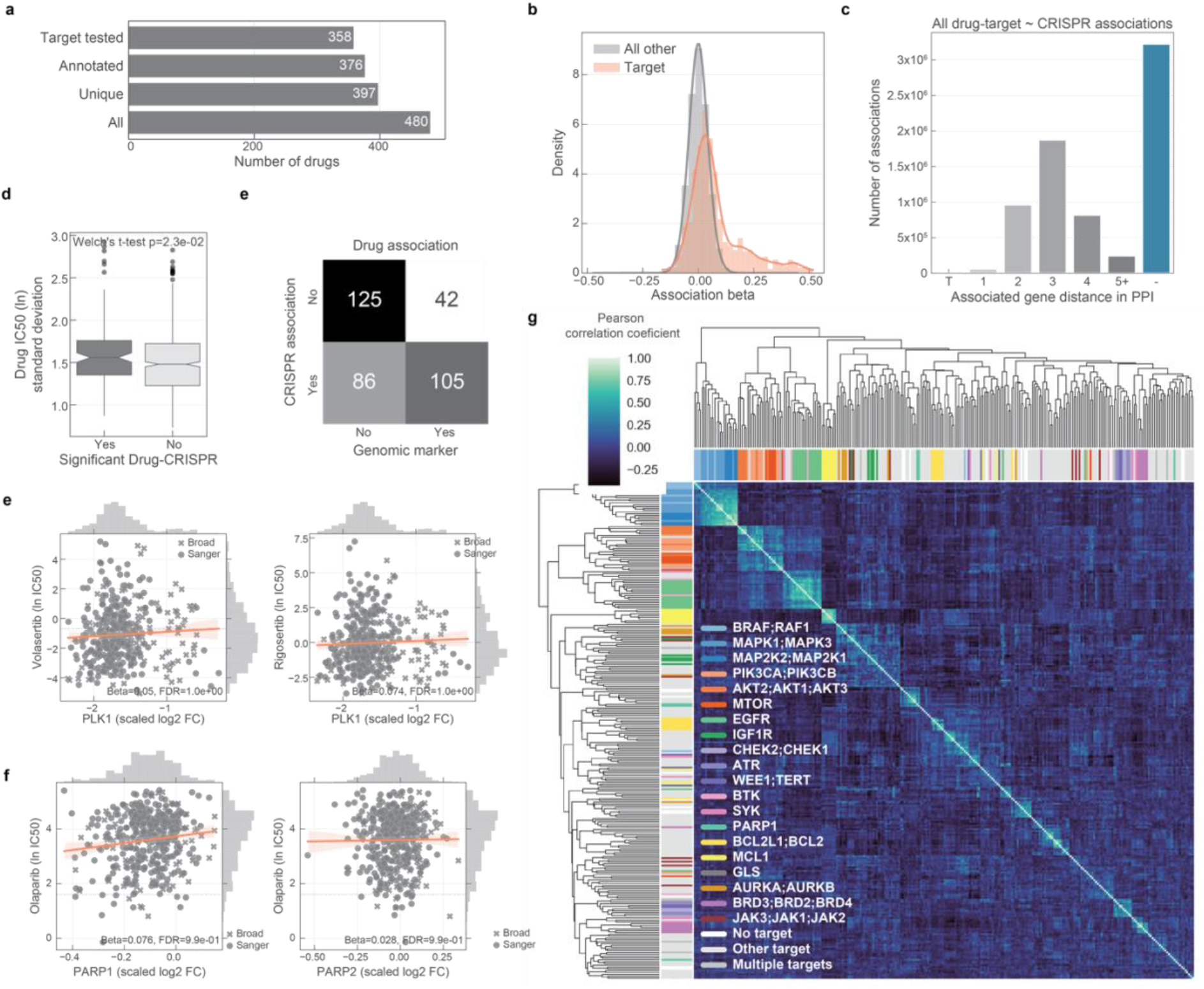
Drug response and gene fitness associations. **a**, total number of drugs utilised in the study and the different levels of information available: ‘All’ represents all the drugs including replicates screened with different technologies (GDSC1 and GDSC2); ‘Unique’ counts the number of unique drug names; ‘Annotated’ shows the number of unique drugs with manual annotation of nominal targets; and ‘Target tested’ represents the number of unique drugs, with target information, for which the target has been knocked-out in the CRISPR-Cas9 screens. **b**, histogram of the drug-gene associations effect sizes (beta) highlighting drug-target associations. **c**, distribution of the shortest path lengths between all the tested drug-gene pairs. For drugs with multiple targets the smallest shortest path of all the targets was taken. **d**, PLK1 inhibitors drug response correlation with PLK1 knockout log2 fold change (FC) gene fitness effects. The dashed grey line indicates the dose response highest drug concentration. **e**, similar to d, correlation of olaparib drug response and both targets PARP1 and PARP2 gene fitness effects. **f**, drug-target associations split by significance (FDR < 10%) plotted against the standard deviation of the drug IC50 (ln) measurements of the respective pair. **g**, contingency matrix of significant drug associations with CRISPR fold changes and binarised event matrix of genomic features, i.e. mutations and copy number gain or loss. **h**, correlation heatmap of the drug-gene effect size across all the genes. Drugs are coloured according to their targets.

**Supplementary Figure 4.**
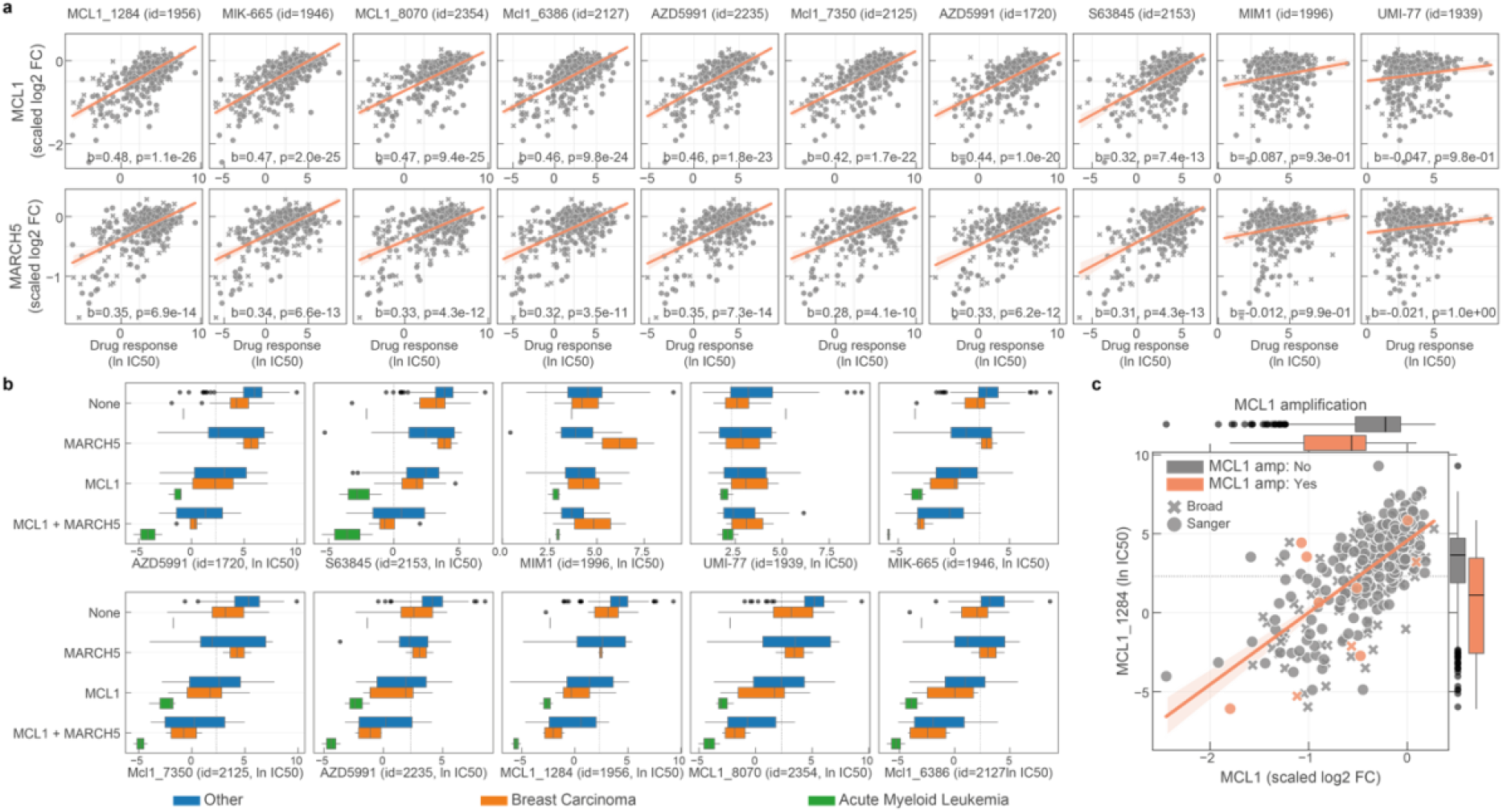
MCL1 inhibitors. **a**, correlation of all MCL1 inhibitor IC50 values against MCL1 and MARCH5 gene fitness profiles. Effect sizes (b) and FDR (p) of the association are reported on the bottom. **b**, stratification of the MCL1 inhibitors drug response measurements according to the cell line dependency on MARCH5 and/or MCL1. Gene vulnerabilities are independent from each other, meaning knockouts were introduced independently and not at the same time. Responses are then split according to the cancer type of the cell lines. Vulnerable cell lines to MARCH5 and MCL1 knockout were defined as those with a depletion of at least 50% of that visible for essential genes (scaled log2 fold change < −0.5). **c**, representative example of a MCL1 inhibitor and their relation with MCL1 gene fitness, with cell lines containing copy number amplification of MCL1 highlighted in orange. Copy number amplified cells were defined taking into consideration their ploidy status, cells with (ploidy <= 2.7 and copy number >= 5) or (ploidy > 2.7 and copy number >= 9) were considered as having MCL1 amplified.

**Supplementary Figure 5.**
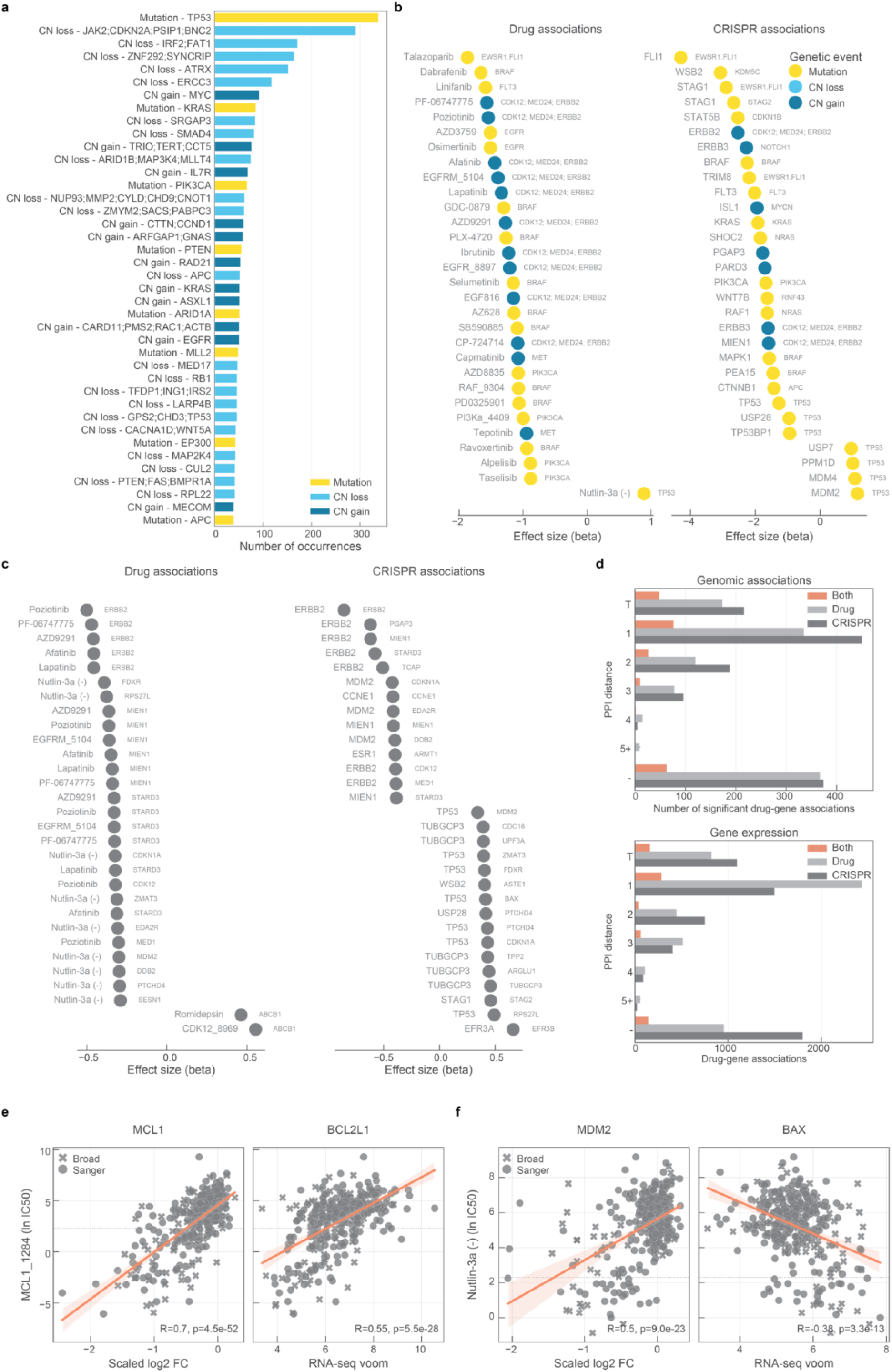
Robust pharmacological associations. **a**, most frequent genomic alterations across the cancer cell lines. Most significant associations between **b**, genomic features and **c**, gene expression profiles with drug response and gene fitness. **d**, number of significant drug-gene pairs across the different types of interactions. Drug-gene pairs were categorised considering the shortest path length between the drug targets and the associated gene. **e**, robust pharmacological association between the expression of BCL2L1 and the significantly correlated pair of MCL1_1284 drug and MCL1 gene fitness profile. **f**, similarly to **e**, but instead it represents a robust pharmacological association between BAX and MDM2 and Nutlin-3a.

## Supplementary Table Legends

**Supplementary Table 1.** Annotated list of the cancer cell lines used in this study.

**Supplementary Table 2.** List of cancer drugs considered in the study.

**Supplementary Table 3.** Drug response matrix (natural log IC50s) for 480 cancer drugs.

**Supplementary Table 4.** Gene fitness CRISPR-Cas9 scaled fold change.

**Supplementary Table 5.** Significant drug-CRISPR associations.

**Supplementary Table 6.** Significant drug-gene expression associations.

**Supplementary Table 7.** Significant robust pharmacogenomic associations with genomic mutation and copy number alterations.

**Supplementary Table 8.** Significant robust pharmacogenomic associations with gene expression.

## Acknowledgements

We would like to thank all members of the Translational Cancer Genomics team who provided useful insight. We are grateful for the diligent contributions of Jon Winter-Holt in terms of compound identification and data clearance and Pedro Beltrao for helpful discussions. Work in M.J.G lab was funded by Wellcome (206194) and AstraZeneca.

## Author contributions

Conceptualization E.G. and M.G.; Formal analysis E.G.; Data curation E.G., C.P., D.v.d.M., A.B., H.L., J.L., B.S., C.C., F.I., S.F. and M.G.; Drug response acquisition and processing D.v.d.M., A.B., H.L. and GDSC Screening Team; Drug annotation E.G., A.S., G.P., F.M.B, P.J., E.C., A.L., C.C. and M.J.G.; Writing original draft preparation E.G. and M.G.; Writing, reviewing and editing all authors; Visualisation E.G.; Supervision: A.L., J.L., B.S., C.C., F.I., S.F. and M.J.G.; Funding acquisition: S.F. and M.J.G.

